# Foveal RGCs develop altered calcium dynamics weeks after photoreceptor ablation

**DOI:** 10.1101/2023.05.30.542908

**Authors:** Zhengyang Xu, Karteek Kunala, Peter Murphy, Laura Patak, Teresa Puthussery, Juliette McGregor

## Abstract

**Objective or purpose:** Physiological changes in retinal ganglion cells (RGCs) have been reported in rodent models of photoreceptor (PR) loss but this has not been investigated in primates. By expressing both a calcium indicator (GCaMP6s) and an optogenetic actuator (ChrimsonR) in foveal RGCs of the macaque, we reactivated RGCs *in vivo* and assessed their response in the weeks and years following PR loss.

**Design:** We used an *in vivo* calcium imaging approach to record optogenetically evoked activity in deafferented RGCs in primate fovea. Cellular scale recordings were made longitudinally over a 10 week period following photoreceptor ablation and compared to responses from RGCs that had lost photoreceptor input more than two years prior.

**Participants:** Three eyes received photoreceptor ablation, OD of a male *Macaca mulatta* (M1), OS of a female *Macaca fascicularis* (M2) and OD of a male *Macaca fascicularis* (M3). Two animals were used for *in vivo* recording, one for histological assessment.

**Methods:** Cones were ablated with an ultrafast laser delivered through an adaptive optics scanning light ophthalmoscope (AOSLO). A 0.5 s pulse of 25Hz 660nm light optogenetically stimulated RGCs, and the resulting GCaMP fluorescence signal was recorded using AOSLO. Measurements were repeated over 10 weeks immediately after PR ablation, at 2.3 years and in control RGCs.

**Main Outcome measures:** The calcium rise time, decay constant and sensitivity index of optogenetic mediated RGC were derived from GCaMP fluorescence recordings from 221 RGCs (Animal M1) and 218 RGCs (Animal M2) *in vivo*.

**Results:** Following photoreceptor ablation, the mean decay constant of the calcium response in RGCs decreased 1.5 fold (1.6±0.5 s to 0.6±0.3 s SD) over the 10 week observation period in subject 1 and 2.1 fold (2.5±0.5 s to 1.2±0.2 s SD) within 8 weeks in subject 2. Calcium rise time and sensitivity index were stable. Optogenetic reactivation remained possible 2.3 years after PR ablation.

**Conclusions:** Altered calcium dynamics developed in primate foveal RGCs in the weeks after photoreceptor ablation. The mean decay constant of optogenetic mediated calcium responses decreased 1.5 - 2-fold. This is the first report of this phenomenon in primate retina and further work is required to understand the role these changes play in cell survival and activity.

## Introduction

In diseases such as age-related macular degeneration and retinitis pigmentosa, photoreceptor death leads to irreversible vision loss. A range of vision restoration therapies are currently under development that aim to restore light sensitivity to the retina including optogenetic and optoelectronic prostheses, and regenerative stem cell transplantation. However, all current vision restoration approaches that act at the retinal level require reactivation of the remaining retinal architecture to restore interpretable signals to the brain. These approaches therefore require not only structural preservation of the inner retina, but high-quality functional preservation following extended periods of vision loss.

Previous studies have demonstrated RGC survival following long-term photoreceptor loss both in mice^1,2^ and primates^3–5^. While RGCs may survive for many years in cases of advanced retinal degeneration, there is evidence of cytopathologies and ultrastructural ‘whorls’ suggestive of organelle stress^6^. Physiological studies of RGCs in murine models of retinal degeneration report increased spontaneous activity, including a higher oscillatory firing frequency and an increased sustained firing rate^7–10^. In primates, structural rewiring of the midget pathway has been reported following acute photoreceptor loss^11^, but physiological changes in primate RGCs have yet to be investigated. If altered physiology does develop in primate RGCs in the absence of photoreceptor input, this may compromise the efficacy of vision restoration therapies. In this study, we investigated the *functional* preservation of RGCs after photoreceptor ablation with the aim of understanding how deafferented RGCs in the fovea respond to restored activity with increasing time after photoreceptor loss.

In humans, high acuity vision is mediated by the specialized anatomy and physiology of the fovea. Previous attempts to restore vision in patients with retinal prostheses have been most successful when the fovea is stimulated^12^, and optimizing therapies at this location may provide a pathway toward improved visual outcomes^13,14^. Due to the degree of similarity in foveal anatomy, physiology, and downstream processing, the non-human primate model is the gold standard animal model for preclinical development of vision restoration therapies. While naturally occurring retinal degeneration models have recently been identified in NHPs^15,16^, they are not widely available at present. Here we adopt a photoreceptor ablation approach using a femtosecond laser exposure delivered through an adaptive optics scanning laser ophthalmoscope (AOSLO). This results in acute photoreceptor loss that is laterally and axially confined^17^.

In the absence of photoreceptors, investigating the functional status of deafferented RGCs requires an alternative stimulation approach. In retinal explants electrical stimulation of RGCs is typical, however longitudinal monitoring of responses in these preparations is impossible. In this study, we take an *in vivo* retinal imaging approach, optically stimulating and recording from deafferented RGCs for weeks, months, and years after photoreceptor ablation by combining AOSLO with the optogenetic actuator ChrimsonR and the calcium indicator GCaMP6s expressed in the foveal RGCs of the NHP. Calcium imaging with AOSLO enables a longitudinal readout of the intracellular calcium response to stimulation at the cellular scale *in vivo*^18^. In this study, we presented a brief pulsed optogenetic stimulus focused onto deafferented RGCs, and recorded the optogenetic mediated calcium dynamics in the weeks and years following photoreceptor loss. This approach allowed us to observe the impact of localized photoreceptor loss on the calcium dynamics of the same group of RGCs over time and in doing so make observations that have previously been experimentally inaccessible.

## Methods

### Animal Care and housing

The primates in this study were housed in an AAALAC-approved facility, where the animals were provided with free access to water and nutritious chow together with various treats and vegetables. Primates were provided with daily enrichment including 2-4 pieces of manipulata such as mirrors and puzzle feeders rotating between animals, daily movies and music, and rotating access to larger free roaming space with swings and elevated perches. A team of 4 full-time veterinarians, 5 veterinary technicians, and an animal behavior specialist monitored animal health, body condition and checked for any signs of discomfort daily. Subjects were housed in diffuse ambient illumination during the daytime (equivalent to 620 nm 0.05 mW at the pupil plane), significantly lower than the ChrimsonR activation threshold^19^, thus deafferented RGCs were only stimulated during the optogenetic trials. This study was performed in strict accordance with the Association for Research in Vision and Ophthalmology (ARVO) statement for use of animals, and the recommendations in the Guide for Care and Use of Laboratory Animals of the National Institutes of Health. All procedures were approved by the University of Rochester Committee in animal resources (PHS Assurance No. 1160743209A1).

### Co-expression of GCaMP6s and td-Tomato labeled ChrimsonR

AAV2-CAG-tdTomato-ChrimsonR and AAV2-CAG-GCaMP6s were synthesized by the University of Pennsylvania vector core and intravitreally injected into the left eye of one female *Macaca fascicularis* (M1) 65 weeks before recording, the right eye of one male (M2) 11 weeks before recording, and the right eye of a male *Macaca fascicularis* (M3) 164 weeks before euthanasia. 1.425e+12 viral genomes of GCaMP6s & 7.88e+10 viral genomes of ChrimsonR were co-injected in subjects M1 and M2 while 1.05e+12 viral genomes of ChrimsonR & 1.94e+13 viral genomes of GCaMP6s were co-injected in subject M3. We co-injected AAV2-CAG-tdTomato-ChrimsonR and AAV2-CAG-GCaMP6s simultaneously to prevent the development of inactivating antibodies to AAV2, which might otherwise preclude expression of the second vector if injections are done sequentially. Simultaneous bilateral injections were never performed. Ocular tissues were sterilized with 50% diluted Betadine prior to the injection and a lid speculum was placed. The viral vector was injected into the middle of the vitreous from a location ∼3 mm behind the limbus using a 30-gauge needle with a tuberculin syringe. Neutralizing antibodies to AAV2 were measured in serum prior to purchase of NHPs and only animals which showed an antibody response to a serum dilution of 1:25 or less were included in the study. Animals received daily subcutaneous cyclosporine A injections to suppress the immune system 1-4 weeks prior to and after injection of the viral vectors. A starting dose of 6 mg/kg was titrated into a therapeutic range of 150–200 ng/mL by monitoring blood trough levels. Once the desired blood trough level was achieved, that dose was maintained throughout the experiment. Signs of intraocular inflammation were monitored by slit lamp examination, fundus photography, and observation of the pupillary response. OCT scans of the primate fovea pre and post AAV injection showed no gross signs of inflammation such as retinal thickening or vitreous cells either immediately after injection or at the time of data collection in any of the three subjects used in this study (Supp. Figure 1).

### Photoreceptor Ablation by ultrafast laser exposure

A patch of photoreceptors was ablated in the central fovea by 106 ms exposure of a 0.8 x 0.7° field to an ultrafast, pulsed, 730 nm laser focused onto the photoreceptor layer (PRL) using the AOSLO (average power of 4.48 Wcm^−2^ and a repetition rate of 80 MHz). Dhakal *et al.* have previously shown that this approach creates damage to the PRL that is both axially and laterally confined^17^. Furthermore, as the RGCs in the fovea are laterally displaced from the photoreceptors that drive them, the deafferented RGCs under study here were never directly exposed to the high intensity laser during PRL ablation. This femtosecond laser ablation approach has previously been used as a model of vision loss in photoreceptor transplantation studies^20^ and to demonstrate optogenetic restoration of RGC activity^5,18^.

### Fundus imaging/OCT/cSLO Imaging

Imaging was performed with a conventional scanning light ophthalmoscope (Heidelberg Spectralis), OCT, and a fundus camera (Topcon TRC 50ex) prior to and weekly following PRL ablation. The fundus camera was equipped with customized fluorescence filters for GCaMP imaging (excitation 466/40 nm, emission 520/28nm) and tdTomato imaging (excitation 549/25 nm and emission 586/20 nm) to assess the fluorescence of GCaMP6s and tdTomato-ChrimsonR independently. The axial impact of laser exposure was monitored with OCT.

### Adaptive optics calcium imaging

Foveal RGCs were stimulated and recorded through an AOSLO system described in Gray *et al*^21^. To summarize, a Shack-Hartmann wavefront sensor and a deformable mirror in the subject pupil plane detected and compensated ocular aberrations in a closed-loop correction system using 847 nm laser diode (Thorlabs) in an SLO. A 796 nm superluminescent-diode laser source (Superlum) was focused on the photoreceptor mosaic layer, and reflectance images collected through the 25.6 Hz adaptive optics SLO with a 2-airy disk, confocal pinhole. GCaMP6s was excited by focusing a 488 nm laser source (Qioptics) at the RGC layer over an area of 398×398 µm^2^ and emitted photons detected in a 517/20 nm band using an 8-airy disk pinhole to maximize signal collection.

### Visual stimulation

Two types of stimuli were delivered through the 25 Hz AOSLO system; a periodic photoreceptor stimulus to assess the loss of photoreceptor function, and a second stimulus to trigger optogenetically mediated calcium responses in RGCs. Both types of trials began with a period of adaptation to the imaging light (67.5 sec for the optogenetic trials and 60 sec for the photoreceptor trials). For the optogenetic stimulus, a 0.5 sec exposure of 25Hz, 640 nm light was focused onto a square region of the RGC layer to trigger optogenetic stimulation, (stimulus mean luminance 21.37 mW/cm^2^, 205×205 µm^2^). Calcium responses were recorded for 22.5 s following stimulation to capture both the rise and decay in response to the optogenetic stimulus. For the periodic photoreceptor stimulus, a 561 nm, (0.5 Hz for M1 & 0.025 Hz for M2) square wave uniform flicker stimulus with a mean luminance of 0.25 mW/cm^2^ was delivered to central photoreceptors for 80 seconds after adaptation while GCaMP recordings were made from the laterally displaced foveal RGCs they drive. In figure 1 a second type of periodic photoreceptor stimulus was delivered using a red 660 nm LED through a Maxwellian view system (0.14 mW/cm^2^, adapted with 488 imaging light for 60s before stimulus was placed).

**Fig. 1:**
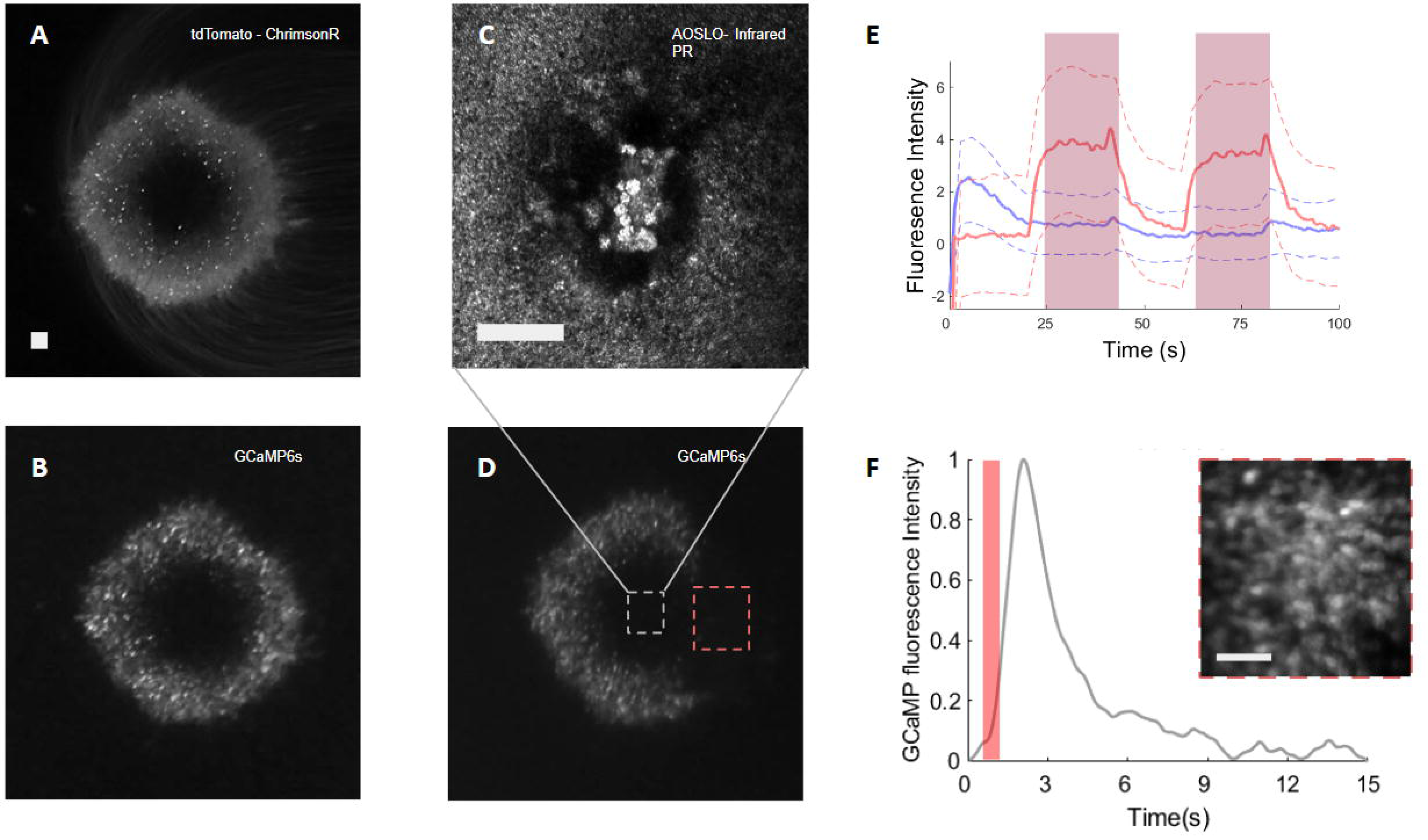
Optogenetic mediated calcium response triggered in deafferented primate foveal RGCs. (A) Fundus fluorescence image of td-Tomato ChrimsonR expression 7 weeks after injection of the viral vector and 1 week prior to photoreceptor ablation. (B) Fundus fluorescence image of GCaMP6s expression 7 weeks after injection of viral vector and 1 week prior to photoreceptor ablation. (C) AOSLO NIR reflectance image of photoreceptor ablation site 1-week following ablation. (D) Fundus fluorescence image of GCaMP nine weeks after injection of viral vector and 1 week post photoreceptor ablation. The dark region of the RGC ring has photoreceptor inputs ablated and the activity dependent fluorescence is correspondingly reduced. (E) A clear 0.025Hz GCaMP response is observed from RGCs on the right side of the RGC ring before photoreceptor ablation, while no activity dependent GCaMP response is observed from the same group of RGCs following photoreceptor ablation, supporting loss of photoreceptor inputs. The traces are an average response collected from 87 deafferented RGCs over a single stimulation period. Response baselines are raised to the same level based on the last 100 frames of the recording for visual comparison. (F) The optogenetic mediated GCaMP response to a 0.5 s 640 nm stimulation 1 week following laser ablation averaged from 87 deafferented RGCs. Stimulus duration is shown by the red bar, AOSLO image of deafferented RGCs with reduced activity dependent fluorescence in top right. All scale bars are 100 μm, and all data were acquired from subject M1.

### Experimental Design

Trials were performed prior to PRL ablation to ensure that we were able to generate robust and reproducible photoreceptor mediated and optogenetic mediated RGC responses. In the 1-5 weeks post ablation, photoreceptor stimulation trials were performed to ensure complete deafferentation of the RGCs under study. To assess changes in the calcium response of RGCs in the weeks following deafferentation, optogenetic stimulation was performed both in the recently deafferented area and in M2, in an eccentricity matched region of RGCs that had been deafferented for over 2 years on the opposite side of the RGC ring. These experiments were repeated over the 10 week period immediately following PRL ablation in both animals. Control experiments were also performed including presentation of the 488 nm excitation light alone to confirm the responses were stimulus dependent and an optogenetic stimulus only trial to ensure no bleed through of the 640 nm stimulus into the detection channel.

### Data analysis

To correct for eye motion *in vivo,* fluorescence videos containing the calcium imaging data from the ganglion cell layer were co-registered to simultaneously acquired, higher signal to noise IR reflectance images of the photoreceptor mosaic. For each imaging location, a reference frame was used to perform a strip-based cross-correlation^22–24^ to remove translation and rotation. Motion corrected reflectance and fluorescence videos were summed over time to generate higher signal to noise reflectance images of the photoreceptor ablation site and fluorescence images of intact and deafferented RGCs for manual segmentation. To ensure the same group of cells were tracked longitudinally, the fluorescence GCaMP images of RGCs from each time point were registered to week 1 post ablation, prior to RGC segmentation.

To extract GCaMP recordings for each RGC, a segmentation mask was applied to the motion-corrected fluorescence videos and the initial 67.5 s adaptation period was removed. The mean signal within each individual cell mask was computed for a single frame (subject M1 96 RGCs, subject M2 109 RGCs).

In optogenetic stimulation trials, each trace was separated into a rising phase and a decay phase based on the frame containing the maximum fluorescence intensity. Rise time was calculated as the time elapsed (number of frames x frame duration) between the stimulus onset and the frame containing the maximum fluorescence intensity. To determine the decay constant for single traces, the data was first normalized by subtracting the average baseline fluorescence in the 100 frames prior to stimulus onset. The resulting traces were fitted with an exponential function using a least-square approach (2 variables, exponential decay constant & magnitude scaler). The mean responses presented in figures 2 and 3 were created by taking the mean of the normalized single cell traces from all segmented RGCs for each location.

**Fig. 2:**
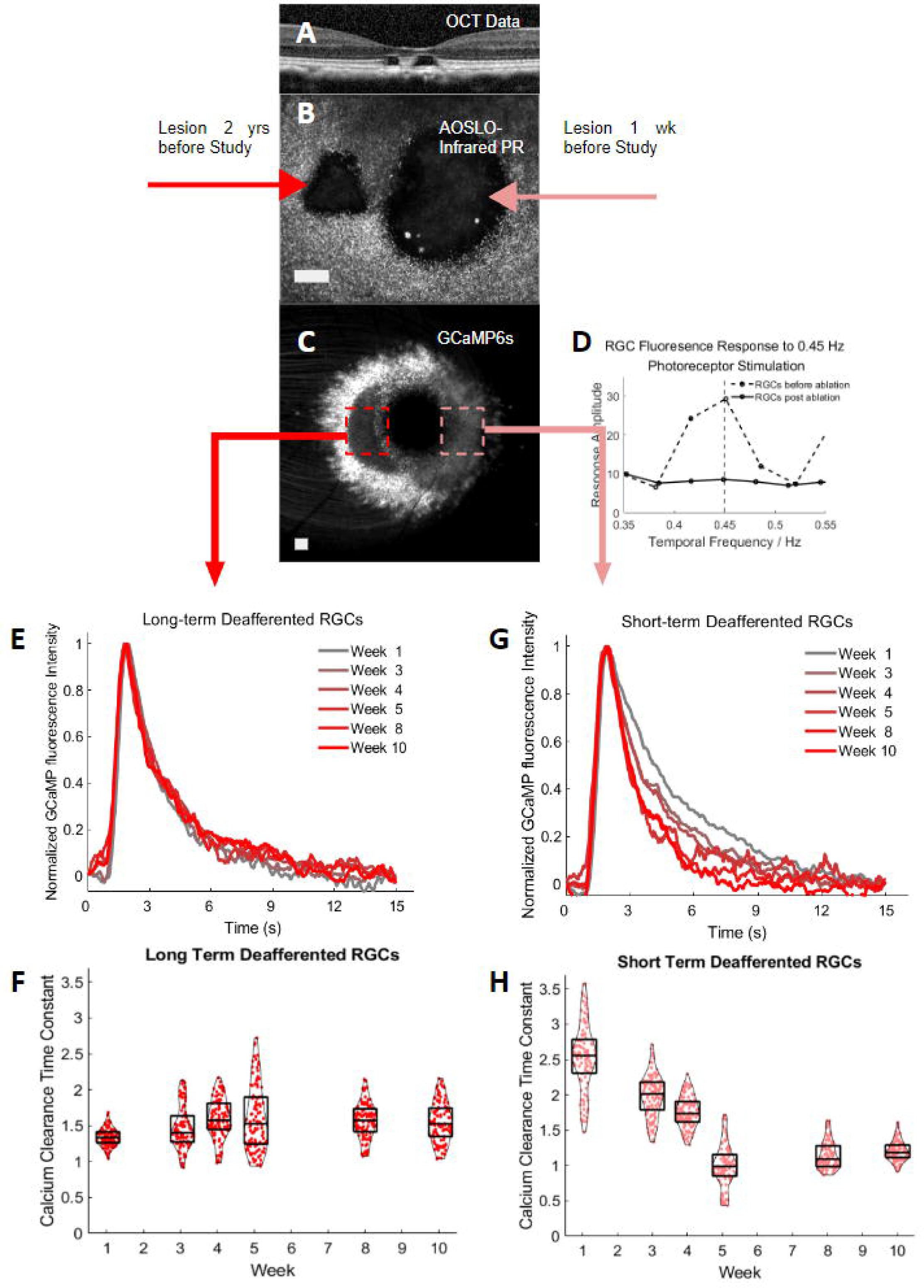
Foveal RGCs show faster calcium decay in the five weeks following PRL ablation in subject M2. (A) OCT scan through the foveal region shows two localized regions of photoreceptor loss. (B) AOSLO NIR reflectance image of two photoreceptor ablation sites 122 weeks following the first ablation (left) and 10 weeks following the second ablation (right). (C) Fundus fluorescence image of GCaMP6s fluorescence 1 week following the second PRL ablation. Two dark regions of the RGC ring have photoreceptor inputs ablated on the nasal and the temporal side of the RGC ring, reflecting the laser PRL ablation performed 2 years and 1 week before the study. The nasal side corresponds to the short-term deafferented RGCs, and the temporal side corresponds to the long-term deafferented RGCs. (D) A 0.45 Hz GCaMP response is observed in RGCs before photoreceptor ablation, while no GCaMP response is observed from the same group of RGCs following photoreceptor ablation. (E) Mean GCaMP response to optogenetic stimulation from 109 long-term deafferented RGCs from weeks 1-10. (F) GCaMP fluorescence decay constants from 109 individual long-term deafferented RGCs from weeks 1-10. (G) Mean GCaMP response to optogenetic stimulation from 109 short-term deafferented RGCs from weeks 1-10. (H) GCaMP fluorescence decay constants from 109 individual short-term deafferented RGCs from weeks 1-10.

The response magnitude was quantified in the following ways:

#### Absolute response magnitude

Calculated to report the response magnitude excluding changes in signal baseline as F_LT_-F0_LT_ & F_ST_-F0_ST._ This is the difference between the maximum fluorescence signal (F) in a single cell and the pre-stimulus baseline (F0) in long term (LT) and short term (ST) deafferented RGCs.

#### Normalized response magnitude

Then the normalized response magnitude is calculated by dividing the absolute response magnitude for each RGC (F_ST_-F0_ST_) by the mean of F_LT_-F0_LT_ recorded in the same imaging session to exclude changes in eye quality or imaging conditions over weeks. The sequence of experiments between sessions was not varied over the sessions.

#### Normalized pre-stimulus baseline

The normalized F0 value was acquired by dividing the short-term deafferented RGC fluorescence baseline for each cell level by the mean of long-term deafferented RGC fluorescence baseline recorded within the same imaging session.

#### The sensitivity index

((F-F0) / std(F0)) was measured by dividing the absolute magnitude of individual deafferented RGC’s F-F0 by the standard deviation of the pre-stimulus baseline. F-F0 & F0 were calculated with the same method described above. The sensitivity index is a modified version of the z-score. The z-score measures the signal magnitude minus the mean, divided by the standard deviation of the entire recording, whereas the sensitivity index measures the signal magnitude minus the mean divided by the standard deviation of the pre-stimulus period.

For photoreceptor stimulation trials, the 60-s adaptation period is removed from the registered video, and the mean signal within each cell mask is computed for each frame of the recording period. The resulting fluorescence traces were multiplied by a Hann windowing function to remove ringing artifacts and Fourier transformed into the frequency domain as described previously^18^.

### Light safety

The total light exposure at each retinal location was calculated for each recording session, with all light sources taken into account (488 nm at 0.71 mw cm^−2^, 796 nm at 1.30 mw cm^−2^, and 843 nm at 0.19 mw cm^−2^), optogenetic stimulation (640 nm at 21.37 mw cm^−2^), and photoreceptor stimulation (561 nm at 0.25 mw cm^−2^). Overall light exposure at a single retinal location did not exceed the American National Standard Institute (American National Standard for the Safe Use of Lasers ANSI Z136.1–2007) maximum permissible exposure of the human retina scaled by the ratio of numerical aperture difference between human and macaque (Photoreceptor ablation session excluded). The power of each light source was measured before the imaging session at the pupil plane, and accumulation of laser exposure between sessions is not considered since all sessions were separated by at least 5 days.

### Histology

#### Tissue preparation

Subject M3 was euthanized with an intravenous injection of Euthasol and perfused with 1 L of heparinized saline for 23 min and 1 L of 4% paraformaldehyde in 0.1M phosphate buffer for 7 min. After enucleation, the anterior segment was removed and the posterior segment was immersion-fixed in 4% paraformaldehyde for an additional 90 min. After fixation, the posterior segment was washed for 5 min in PBS and cryoprotected in 10%, 20%, and 30% sucrose. Retinal pieces were embedded in Cryogel embedding compound (Leica) and frozen in isopentane cooled in liquid nitrogen. 20-μm thick cryosections were collected and dried for 60 min at 35°C on a slide warmer before storage at −20°C. Sections were frozen at −20°C until further use.

#### Immunostaining

To assess the nuclear to cytoplasmic ratio of GCaMP6s expression in RGCs, nuclei were counterstained with Hoescht 33342. To assess the retinal cell types that were transduced by GCaMP6s/tdTomato, retinal sections were immunostained with an RBPMS antibody (RRID: AB_2492226, Phosphosolutions, Cat #1832. 1:200) to label all RGCs^25^ and a CAVIII antibody (RRID: AB_2066293, Santa Cruz Biotech, SC-67330, 1:50) to label parasol RGCs^26,27^. After a 5-minute wash in PBS, retinal sections were blocked for 1hr in 10% normal horse serum (NHS), 1% Triton X-100, 0.025% NaN_3_ in PBS. The primary antibodies were incubated overnight at room temperature diluted in 3% NHS, 1% Triton X-100, 0.025% NaN_3_ in PBS. Retinal sections were then washed in PBS and incubated in secondary antibodies diluted in 3% NHS, 0.025% NaN_3_ in PBS for 1 hr at 22C. The secondary antibodies were Donkey anti-Rabbit Alexa 594, RRID: AB_141637, Molecular Probes: A-21207, 1:800 and Donkey anti-Guinea Pig Alexa 647 (RRID: AB_2340476, Jackson ImmunoResearch: 706-605-148, 1:500). After washes, nuclei were counterstained with Hoescht 33342 then mounted in Mowviol.

#### Confocal imaging

Samples were imaged on a laser scanning confocal microscope (Zeiss LSM 880) with the 405-, 488-, 561-, 594-, and 633-laser lines using a Zeiss Plan-Apochromat 60×/1.4 N.A. oil immersion or a Zeiss Plan-Apochromat 20x/0.8 air objective at 1024 x 1024 resolution. Tiled image stacks were stitched in Image J or Zen Blue software (Zeiss).

#### Analysis

To measure the nuclear to cytoplasm ratio (N/C) for GCaMP6s fluorescence intensity, deafferented and non-deafferented regions of the GCL were imaged with a 60×/1.4 N.A objective. 20 pixel thick line profiles were drawn through the center of each GCaMP6s positive RGC using Image J to generate intensity profile plots of GCaMP6s and Hoescht fluorescence. Intensity plots were exported to Igor Pro 9 (Wavemetrics) for further analysis. Intensity measurements were background subtracted using regions of the image without GCaMP6s fluorescence. To quantify the percentage of GCaMP6s or tdTomato expressing RGCs that were parasol cells, we counted the number of immunolabeled CAVIII+ RGCs (RGC identity confirmed by RBPMS+ immunolabeling) that also had endogenous GCaMP or tdTomato expression. Quantifications were performed on confocal maximum intensity projections of 5 x 0.7 µm z steps. N/C ratio and transduction quantifications were each performed on n=2 retinal sections from animal M3.

## Results

### Localized photoreceptor ablation in primate fovea deafferents RGCs allowing characterization of optogenetic mediated calcium dynamics

Intravitreal co-injection of the viral vectors AAV2-CAG-GCaMP6s and AAV2-CAG-tdTomato-ChrimsonR resulted in a foveal ring of RGCs expressing the calcium indicator GCaMP6s (Figure 1A and Supp. Figure 2A-C) and the optogenetic actuator tdTomato-ChrimsonR in all 3 animals (Supp. Figure 2D-F) as observed in previous studies^5,18^. We expect that the transduced cells are predominantly midget RGCs given the high proportion of these cells in the fovea (∼80-95%)^28,29^, but we also saw evidence of transduction of some parasol RGCs (Supp. Figure 3).

AOSLO NIR reflectance imaging of the cone photoreceptor mosaic *in vivo* confirmed photoreceptor ablation following laser exposure (Figure 1C). Optical coherence tomography (OCT) showed the absence of the foveal photoreceptor layer (PRL) post exposure (Supp. Figure 1A-C). Figure 1D shows GCaMP fluorescence from foveal RGCs one week after PRL ablation (9 weeks after the initial injection of AAV2-CAG-GCaMP6s and AAV2-CAG-ChrimsonR-tdTomato). As previously reported^5,18^ fluorescence fundus imaging shows a reduction in activity dependent fluorescence from the RGC ring that is spatially consistent with the ablation of photoreceptors in the temporal hemifield of the central fovea. The loss of photoreceptor response was confirmed by presenting a 0.025 Hz square wave flicker stimulus to the central fovea. Figure 1E shows 0.025 Hz GCaMP responses from RGCs prior to laser exposure and the absence of these signals from the same RGCs post exposure as anticipated.

To assess calcium dynamics in these deafferented RGCs we presented a 0.5s, 640 nm, 0.9 mW stimulus focused at the ganglion cell layer through AOSLO to optogenetically stimulate the RGCs. The optogenetic mediated calcium response leads to an increase in activity dependent GCaMP fluorescence and then gradual decay. Figure 1F shows the calcium responses recorded from deafferented RGCs 1 week after PRL ablation including an inset showing an *in vivo* fluorescence image of the RGCs under study.

### Calcium responses to optogenetic stimulation decay more rapidly in RGCs within weeks of PRL ablation

The same laser ablation, optogenetic stimulation and GCaMP recording paradigm was also used to interrogate the calcium rise and decay in a second animal (M2). In this animal, recordings were made from both a patch of RGCs that had been deafferented for 1 week (nasal: short-term deafferented), and also from an eccentricity-matched control patch of RGCs deafferented 2 years previously (temporal: long-term deafferented). By recording from these two regions during each session we were able to use long-term deafferented RGC responses as a control group to assess the stability of the imaging platform.

The NIR reflectance AOLSO image of the photoreceptor mosaic and OCT data show the two regions of photoreceptor ablation (Figure 2A-B) in subject M2. The decreased activity dependent RGC GCaMP fluorescence in Figure 2C reflects the loss of photoreceptor input, consistent with the previous study^5^. Deafferentation was confirmed at the 1-week timepoint post ablation on the nasal side of the RGC ring, with no response to 0.45 Hz photoreceptor stimulation detected (Figure 2D). The rise time and decay constant in the long term deafferented RGCs (101 to 122 weeks post ablation) was stable (Figure 2E-F). By contrast, in the recently deafferented RGCs (Figure 2G-H the mean decay constant of the calcium response decreased 2.1-fold (from 2.5±0.5 s to 1.2±0.2 s SD) in the 8 weeks following PR ablation (*p*<0.001, paired *t*-test, n=109), with 87% of this decrease occurring within the first 5 weeks. The decay constant did not decrease further from week 8 to 10 and approached the value for the long term deafferented RGCs.

### The rate of calcium decay is stable in RGCs with intact photoreceptors and in RGCs prior to PRL ablation

Longitudinal recording from deafferented RGCs was repeated in a second subject (M1) this time comparing the optogenetic response in deafferented RGCs to the optogenetic response in RGCs with intact photoreceptors. Photoreceptor-mediated calcium responses to a 0.1 Hz flickering stimulus incident on foveal photoreceptors were recorded 1 week prior to, and 5 weeks post PRL ablation in a single field of view (Figure 3A-B). Laser ablation in subject M1 removed photoreceptor input from half of the RGCs in the recording area, leaving those at higher eccentricity with inputs intact. This scenario is shown in Figure 3C. After ablation, periodic RGC responses to photoreceptor stimulation were absent in the deafferented zone (Figure 3D-E). Similar to animal M2, the mean decay constant of the optogenetically-evoked calcium responses in the deafferented RGCs decreased 1.5-fold (from 1.6±0.5 s to 0.6±0.3 s SD) in the 10 weeks post PR ablation (*p*<0.001, paired *t*-test, Figure 3F-G). In comparison, RGCs with photoreceptors intact in the same field of view showed relatively stable decay kinetics over the 10-week period. Use of a spatially limited optogenetic stimulus in the fovea ensured that the calcium responses from these non-deafferented cells were not photoreceptor mediated (Supp. Figure 4). Optogenetic stimulation of RGCs with intact photoreceptors in this region and in the deafferented region 1 week prior to photoreceptor ablation produced a stable decay constant (2.2±0.5 s), confirming that these results are not explained by a temporary increase in decay constant relating to e.g. inflammation post ablation (Figure 3H).

**Fig. 3:**
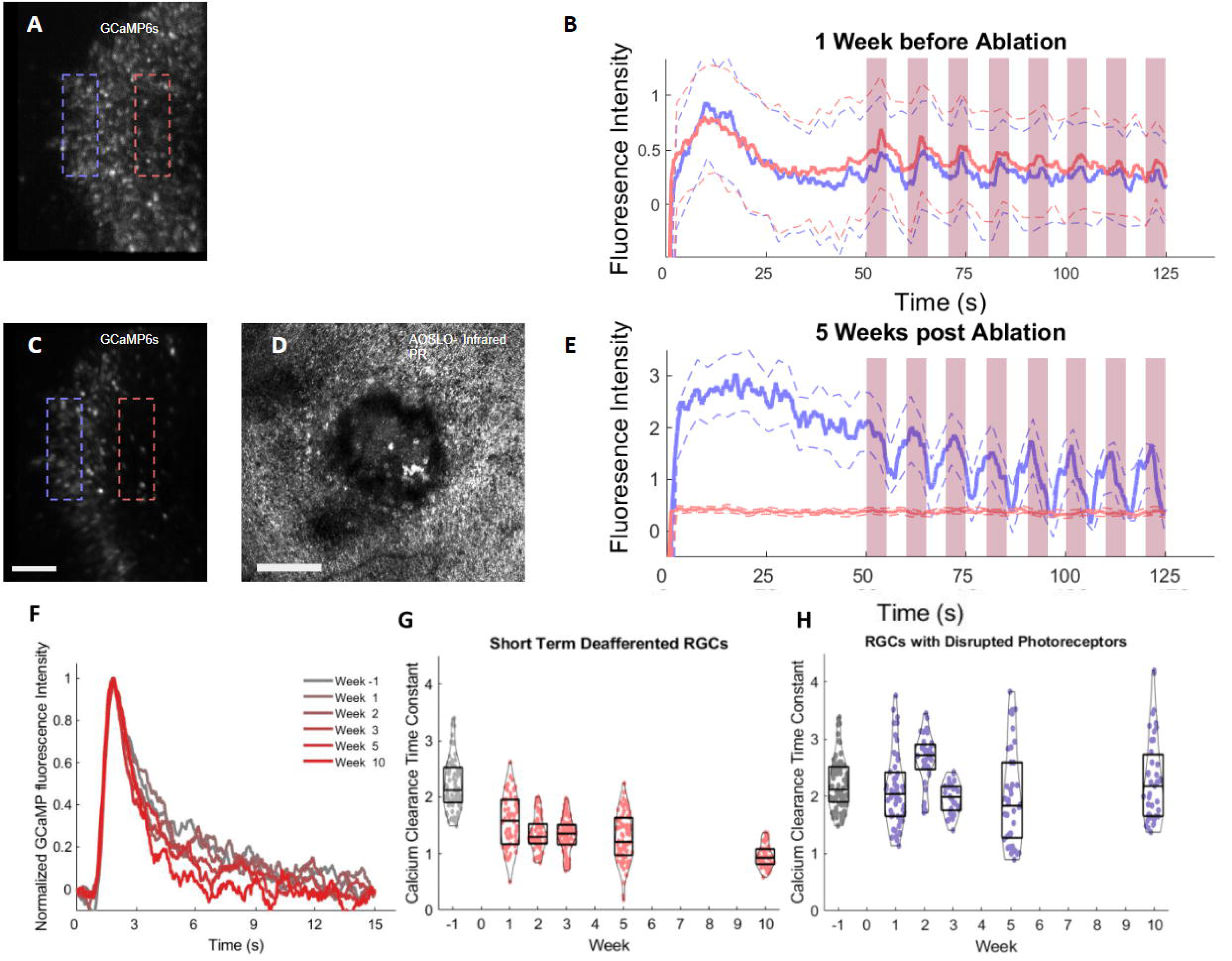
Optogenetic mediated calcium decay over time in deafferented and intact primate foveal RGCs pre and post photoreceptor ablation in a second primate. (A) SLO GCaMP fluorescence image showing the GCaMP fluorescence from the two test regions of foveal RGCs prior to photoreceptor ablation. (B) Photoreceptor mediated GCaMP responses in RGCs to 0.1 Hz flicker stimulation from the purple and pink regions 1 week prior to photoreceptor ablation the traces show the averaged responses of 69 RGCs in the two areas to a single stimulation trial. Response baselines are raised to the same level based on the last 100 frames of the recording for visual comparison, and the stimulus record is shown by the red bar. (C) Same region as shown in (A) 5 weeks post PRL ablation deafferenting the RGCs in the pink region. (D) AOSLO NIR reflectance image of the photoreceptor ablation site 5 weeks after PRL ablation. (E) Photoreceptor mediated GCaMP responses to 0.1 Hz flicker stimulation in deafferented (purple) and intact (pink) areas 5 weeks post photoreceptor ablation, the traces show the averaged responses of 69 RGCs in the two areas to a single stimulation trial. Response baselines are raised to the same level based on the last 100 frames of the recording for visual comparison, and the stimulus record is shown by the red bar. (F) The mean normalized optogenetic mediated GCaMP response recorded from 129 short-term deafferented RGCs over the observation period from 1 week before PRL ablation to 10 weeks post-ablation. (G) Calcium decay constants recorded from 129 single RGCs over the observation period from 1 week before PRL ablation to 10 weeks post-ablation. (H) Simultaneously acquired calcium decay constants recorded from 92 RGCs over the observation period from 1 week before PRL ablation to 10 weeks post-ablation. All scale bars are 100 μm, and all data were acquired from subject M1.

### The optogenetic mediated calcium rise time is unaffected by deafferentation

In animal M1, the mean rise time did not change significantly prior to and post PRL ablation (Week −1, 1.47±0.33s SD) & (Week 5, 1.44±0.14s SD). In animal M2, the mean rise time also did not differ significantly between the long-term (1.43±0.17s SD) and the short-term (1.45±0.30s SD) deafferented RGCs during the 10-week period (unpaired *t*-test, *p*=0.05, n=654) (Figure 4A-B). These data confirm that photoreceptor ablation does not alter the rise time of calcium responses in downstream RGCs and is as expected as the calcium rise time primarily reflects the speed of the calcium indicator GCaMP6s, (optogenetic triggered spiking would be expected to occur on a timescale of milliseconds). While this is less biologically relevant to the optogenetic response, it does highlight the stability of our recording platform.

**Fig. 4:**
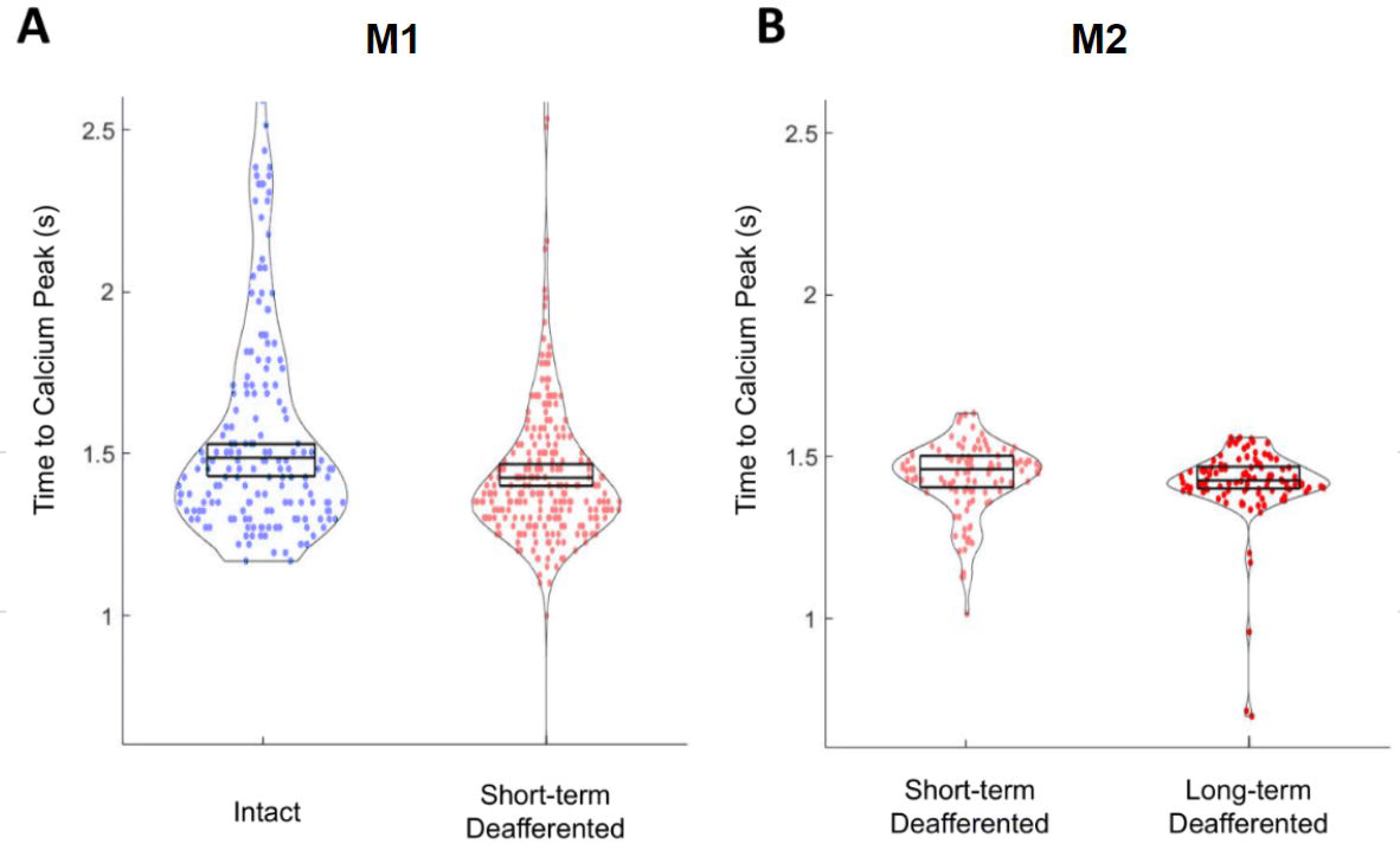
Optogenetic mediated calcium rise time in deafferented and intact RGCs. (A) Optogenetic mediated calcium rise time in response to a 0.5 s, 25 Hz 640 nm stimulus delivered to RGCs with PR input intact (n=182) and RGCs with PR inputs ablated 1-10 weeks previously (short term deafferented) RGCs (n=257) in subject M1. Standard deviations of the RGC rise time are labeled with the upper and lower edge of the box plot, and the mean of RGC rise times are labeled with middle line in the box plot. Each data point shows the rise time recorded from a single RGC within an imaging session. (B) Optogenetic mediated calcium rise time for short term deafferented RGCs (n=106) compared to RGCs deafferented for 2 years (long-term, n=107), in animal M2. Each data point shows the mean RGC rise time collected from a single cell over the six imaging sessions within the 10-week observation period.

**Fig. 5:**
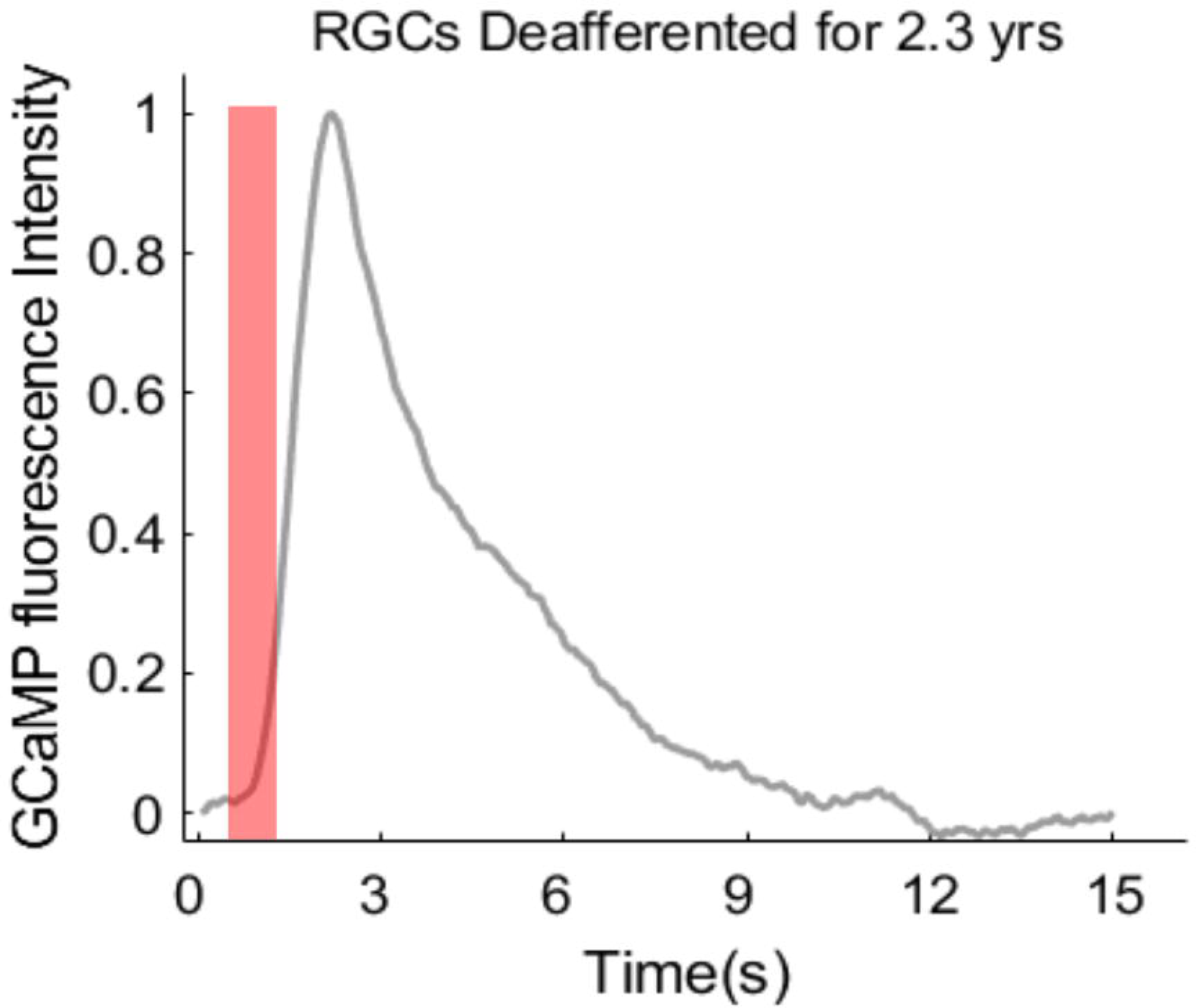
Mean optogenetic mediated calcium response recorded from 109 RGCs deafferented over 2.3 years. Optogenetic mediated GCaMP response to a 0.5 second 25 Hz 640 nm stimulus 2.3 years following photoreceptor ablation and 2.8 years following initial vector injection, the stimulation duration is shown by the red bar.

### An optogenetic mediated calcium response can be evoked in deafferented RGCs 2.8 years following viral transduction and 2.3 years after photoreceptor ablation

In subject M2, robust optogenetic mediated responses were recorded 2.8 years post-injection of the viral vector carrying the ChrimsonR transgene. The RGCs in this area had been deafferented for over 2.3 years at the time of recording. The decay constant in these long-term deafferented RGCs was 1.6 ± 0.2s S.D.

### The decay constant is independent of response amplitude, stimulus power, and chronic expression of GCaMP6s

To assess the impact of potentially varying optical parameters, response magnitude and protein expression on the calcium decay constant, we calculated the sensitivity index of responses and the normalized response magnitude to control for these parameters. In addition, we performed control experiments using a range of optogenetic stimulus powers and conducted a histological investigation of intact and deafferented RGCs in subject M3 with chronic GCaMP6s expression.

The sensitivity index scales the absolute response magnitude by the standard deviation of the baseline fluorescence, providing a discriminability metric for the response amplitude independent of fluorescence intensity. The division controls for parameters that may potentially vary between sessions such as the clarity of the anterior optics, system alignment, AO performance, pinhole position and the level of GCaMP expression. The sensitivity index of responses in deafferented RGCs was stable during the 10-week recording period, suggesting changes in the magnitude of the response do not account for the reported reduction in the decay constant (Fig. 6A).

**Fig. 6:**
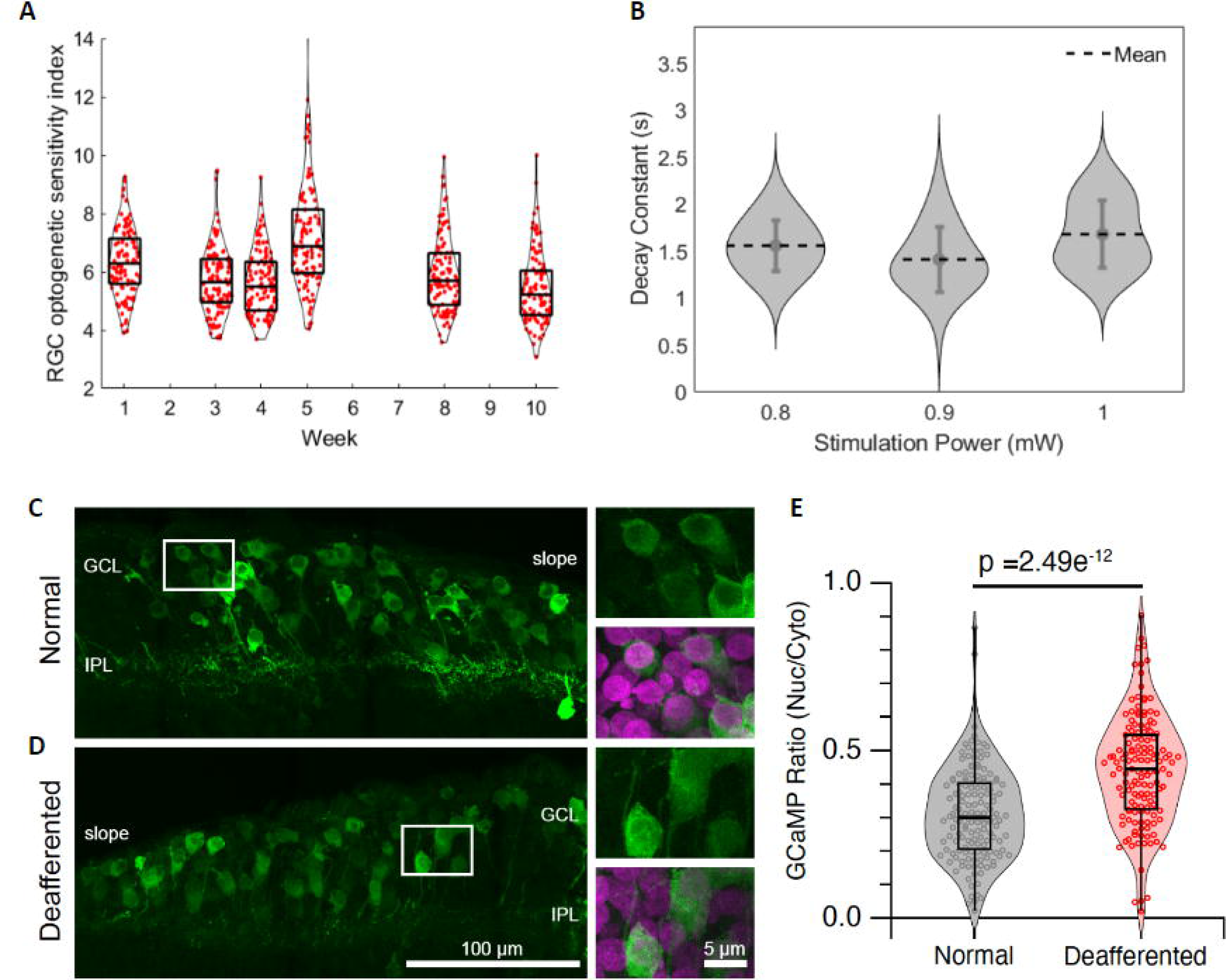
Control experiments showed that the altered calcium dynamics are not explained by changes in response magnitude, stimulus power or nuclear expression of GCaMP6s. (A) Optogenetic sensitivity index of responses to optogenetic stimulation in short-term deafferented RGCs in the 10 weeks following photoreceptor ablation. (B) Calcium decay constant in deafferented RGCs following optogenetic stimulation by three different light intensities. Data collected from 109 short-term deafferented RGCs and 109 long-term deafferented RGCs in subject M2. (C-D), Confocal image (maximal z-projection) showing GCaMP6s expression (green) in a non-deafferented (normal, C) and deafferented (D) region of the same retina from M3. The foveal slope region is indicated. ROIs are shown enlarged on the right with and without a nuclear counterstain (Hoescht 33342). E, Violin/bar plots showing the nuclear:cytosolic ratio of GCaMP6s for RGCs in normal (n=152 cells) and deafferented (n=140 cells) retinal regions from subject 3. The horizontal line shows median, box shows upper and lower quartiles and whiskers show min/max. Dots show data from individual cells. Data were pooled from two different sections of the same retina. Statistical comparison was with an unpaired, two-tailed t-test. GCL, ganglion cell layer, IPL, inner plexiform layer.

Figure 6B shows the decay constant from deafferented RGCs under optogenetic stimulation powers ranging from 0.8 - 1 mW. The decay constants in deafferented RGCs show no linear trend with stimulus power, suggesting the changes we observed in the speed of the calcium decay are not linked to the power of the stimulus or to changes in the level of optogenetic actuator expression over time.

Between week 2 & 3 post PRL ablation, the normalized pre-stimulus GCaMP fluorescence baseline decreased 1.7-fold, consistent with the reduction observed in the fluorescence fundus images (Fig. 2C, Figure S2B). This decrease in baseline fluorescence has been observed and reported in previously work^5^. The normalized response magnitude in the short-term deafferented RGCs also decreased significantly between weeks 1 & 2 following photoreceptor loss (0.11±0.02s to 0.04±0.01s, Figure S2B). The trend of both pre-stimulus baseline and normalized response magnitude change did not follow the gradual decrease in decay constant decrease observed over 5 weeks.

Previous studies in mouse cortical neurons have shown that long-term expression of GCaMP3 can lead to nuclear accumulation and prolongation of the decay of calcium signals^25^. However, recent longitudinal studies with GCaMP7s in rodents showed no abnormal light-evoked activity in RGCs following long-term expression of GCaMP7s^30^. To investigate whether changes in GCaMP6s localization could account for the observed changes in calcium dynamics, we quantified the nuclear to cytosolic (N/C) ratio of GCaMP6s in deafferented and intact RGCs in subject M3 (Figure 6C-E). In prior studies, a higher GCaMP6s N/C ratio was linked to slower decay of calcium signals. We found a significantly higher N/C ratio in deafferented compared to intact RGCs (0.44 ± 0.16 vs 0.31 ± 0.14 S.D.). These findings suggest that the faster decay constant of calcium signals in deafferented RGCs cannot be explained by nuclear accumulation in intact RGCs associated with GCaMP toxicity.

## Discussion

By co-expressing the calcium indicator GCaMP6s and the optogenetic actuator ChrimsonR *in vivo*, we have demonstrated the development of altered calcium dynamics in adult primate foveal RGCs just weeks after photoreceptor loss. The timescale over which these changes were observed is similar to that associated with the development of retinal hyperactivity in rodent models^7–10^. A 1.5 to 2-fold increase in the speed of calcium decay was observed within 5 weeks in subjects M1 and M2 respectively. The decay constant in short-term deafferented RGCs recorded from subject M2 also approached the long-term deafferented RGC responses after week 8. Data from both subjects suggest that physiological changes occurred within weeks of photoreceptor loss. While there was an initial difference in calcium decay constant between the two subjects, it is important to note that both deafferented RGC groups approach a similar decay constant by 10 weeks after photoreceptor loss. The initial difference between the two subjects may be related to the substantial age difference between these animals (M1 7 years old, M2 17 years old, M3 9 years old). Data from post-mortem human retina has suggested that exposure to age-related stressors may generate differences in physiology^6^.

We chose to characterize the calcium dynamics in RGCs using the calcium decay constant, and optogenetic sensitivity index. The decay constant reflects the speed of intracellular calcium clearance from the cell and is similar to the paradigm described by Shiga *et al.*^34,35^, aiming to investigate abnormal calcium clearance in a murine glaucoma model. These authors were ultimately able to connect a lengthening of the calcium decay constant to changes in the endoplasmic reticulum (ER) Ca^2+^ ATPase 2 (SERCA2), which is responsible for pumping Ca^2+^ from the cytoplasm to the ER. As intracellular calcium dynamics are complex, in this study we do not makes claims as to the biological origin of the effect we observe, nevertheless the calcium decay constant remains a tractable metric for initial assessment of changes in calcium clearance occurring in RGCs.

The absolute response magnitude is affected by factors including variable image quality from session to session. To control for these effects, we suggest that the ‘sensitivity index’ which scales the maximum response amplitude by the standard deviation of the signal in the pre-stimulus period is perhaps the most biologically relevant measure of response magnitude as it represents the discriminability of the calcium signal induced by optogenetic stimulation relative to the noise. The normalized response magnitude shows a significant reduction within 1 week following photoreceptor ablation, consistent with the reduction of GCaMP6s and tdTomato-ChrimsonR fluorescence intensity in deafferented RGCs in the weeks of PRL ablation reported by McGregor *et al..* However, as described in the results, the stability of the optogenetic sensitivity index, the stimulus power control experiments, and histological evaluation of GCaMP6s expression suggest that the gradual changes in the decay constant we observe cannot be solely attributed to changes in absolute response magnitude or the level of GCaMP expression but rather represent a biological change in RGCs.

Similarly, the stable decay constant recorded from intact RGCs in subject M1 indicates the altered calcium dynamics observed in both subjects following deafferentation was not a result of optogenetic stimulation per se or a transient response to laser injury. In supplementary figure 4, we also demonstrated that scattered light from the optogenetic stimulation of RGCs did not trigger responses mediated by nearby photoreceptors, confirming the initial longer decay constant observed prior to PRL ablation is optogenetic in origin.

While this is the first report of altered calcium dynamics in primate retina, studies in a murine model of glaucoma have also revealed abnormal calcium dynamics developing in RGCs following axon crush, although interestingly the decay constant increases after injury^34,35^. This is different from the more rapid decay observed in this study, but there are a number of important model and species differences to note. In the glaucoma model in mouse, lengthening of the decay constant appears to precede rapid cell death^34^, whereas in the photoreceptor ablation model in primate the inner retina remains structurally intact despite many years of photoreceptor loss. It is possible that the shorter decay constant we observe is the consequence of a protective adaptation to higher levels of intracellular calcium generated by hyperactivity. Studies in other parts of the primate visual system have observed abnormal calcium homeostasis following ablation^36^. It is also possible that the altered calcium clearance develops following photoreceptor loss to maintain calcium homeostasis. It is also important to note that the results presented here are likely to be dominated by responses from the midget retinal ganglion cell class present only in primate fovea. The inner limiting membrane of the primate retina prevents the direct penetration of the AAV virus from the vitreous into the GCL except through the foveal pit and as a result the RGCs investigated in this study were predominantly midget ganglion cells, with a minor contribution from parasol and other RGC types. Our data show a small fraction of the total cells expressing both GCaMP6s and tdTomato:ChrimsonR. are parasol RGCs (Supp. Fig. 3).

Although physiological changes are observed in RGC calcium dynamics following photoreceptor loss, it is important to note that, robust optogenetic mediated responses were triggered in RGCs even 2.3 years following photoreceptor loss, and 2.8 years after viral transduction of the optogenetic actuator. This is a promising result with current clinical trials having reported optogenetic mediated light sensitivity remaining 1.6 years after administration of the therapy in patients^37^. To fully investigate the impact of altered calcium dynamics on restored vision, psychophysical tests will be necessary.

## Acknowledgements

We thank Bill Merigan and David Williams for accommodating access to the 1P and 2P AOSLOs and sharing animal subjects. We thank Jacqueline Gayet for technical assistance with histology. We thank the Penn Vector Core (RRID: SCR_022432) at the Perelman School of Medicine, University of Pennsylvania This work was supported by the National Institutes of Health National Eye Institute through grants U24 EY033275 (to J.E.M.), EY024265 (to T.P.), P30 Core Grants EY001319 (University of Rochester), EY003176 (UC Berkeley) and T32 EY007043 (L.P.). Additional funding was provided by the Steven E. Feldon Scholarship to J.E.M. and through an unrestricted grant to the Flaum Eye Institute from Research to Prevent Blindness (https://www.rpbusa.org/rpb/grants-and-research/grants/rpb-unrestricted-grants/rpb-unrestricted-grants/). Confocal imaging was conducted in the CRL Molecular Imaging Center at UC Berkeley (RRID:SCR_017852), supported by the Helen Wills Neuroscience Institute.

## References

1. Margolis DJ, Newkirk G, Euler T, Detwiler PB. Functional Stability of Retinal Ganglion Cells after Degeneration-Induced Changes in Synaptic Input. J Neurosci 2008;28:6526.

2. Mazzoni F, Novelli E, Strettoi E. Retinal Ganglion Cells Survive and Maintain Normal Dendritic Morphology in a Mouse Model of Inherited Photoreceptor Degeneration. J Neurosci 2008;28:14282.

3. Santos A, Humayun MS, de Juan E Jr, et al. Preservation of the Inner Retina in Retinitis Pigmentosa: A Morphometric Analysis. Archives of Ophthalmology 1997;115:511–515.

4. Jones BW, Pfeiffer RL, Ferrell WD, et al. Retinal remodeling in human retinitis pigmentosa. Experimental Eye Research 2016;150:149–165.

5. McGregor JE, Kunala K, Xu Z, et al. Optogenetic therapy restores retinal activity in primate for at least a year following photoreceptor ablation. Molecular Therapy 2022;30:1315–1328.

6. Pfeiffer RL, Marc RE, Jones BW. Persistent remodeling and neurodegeneration in late-stage retinal degeneration. Progress in Retinal and Eye Research 2020;74:100771.

7. Stasheff SF. Emergence of Sustained Spontaneous Hyperactivity and Temporary Preservation of off Responses in Ganglion Cells of the Retinal Degeneration (rd1) Mouse. Journal of Neurophysiology 2008;99:1408–1421.

8. Kolomiets B, Dubus E, Simonutti M, et al. Late histological and functional changes in the P23H rat retina after photoreceptor loss. Neurobiology of Disease 2010;38:47–58.

9. Telias M, Denlinger B, Helft Z, et al. Retinoic Acid Induces Hyperactivity, and Blocking Its Receptor Unmasks Light Responses and Augments Vision in Retinal Degeneration. Neuron 2019;102:574–586.e5.

10. Denlinger B, Helft Z, Telias M, et al. Local photoreceptor degeneration causes local pathophysiological remodeling of retinal neurons. JCI Insight 2020;5:132114.

11. Akiba R, Boniec S, Uyama H, et al. Rewiring of the adult midget pathway in a primate model of acute photoreceptor loss. Investigative Ophthalmology & Visual Science 2022;63:3132–3132.

12. Stingl K, Bartz-Schmidt KU, Besch D, et al. Subretinal Visual Implant Alpha IMS – Clinical trial interim report. Vision Research 2015;111:149–160.

13. Damle S, Lo Y-H, Freeman WR. High Visual Acuity Retinal Prosthesis: Understanding Limitations and Advancements Toward Functional Prosthetic Vision. RETINA 2017;37.

14. McGregor JE, Williams DR, Merigan WH. Functional Assessment of Vision Restoration. In: Bowes Rickman C, Grimm C, Anderson RE, et al., eds. Retinal Degenerative Diseases. Cham: Springer International Publishing; 2019:145–149.

15. Moshiri A, Chen R, Kim S, et al. A nonhuman primate model of inherited retinal disease. J Clin Invest 2019;129:863–874.

16. Peterson SM, McGill TJ, Puthussery T, et al. Bardet-Biedl Syndrome in rhesus macaques: A nonhuman primate model of retinitis pigmentosa. Experimental Eye Research 2019;189:107825.

17. Dhakal KR, Walters S, McGregor JE, et al. Localized Photoreceptor Ablation Using Femtosecond Pulses Focused With Adaptive Optics. Translational Vision Science & Technology 2020;9:16–16.

18. McGregor JE, Godat T, Dhakal KR, et al. Optogenetic restoration of retinal ganglion cell activity in the living primate. Nature Communications 2020;11:1703.

19. Klapoetke NC, Murata Y, Kim SS, et al. Independent optical excitation of distinct neural populations. Nature Methods 2014;11:338–346.

20. Aboualizadeh E, Phillips MJ, McGregor JE, et al. Imaging Transplanted Photoreceptors in Living Nonhuman Primates with Single-Cell Resolution. Stem Cell Reports 2020;15:482–497.

21. Gray D. High resolution in vivo imaging of retinal cells with adaptive optics fluorescence scanning laser ophthalmoscopy. 2007.

22. Yang Q, Zhang J, Nozato K, et al. Closed-loop optical stabilization and digital image registration in adaptive optics scanning light ophthalmoscopy. Biomed Opt Express 2014;5:3174– 3191.

23. McGregor JE, Yin L, Yang Q, et al. Functional architecture of the foveola revealed in the living primate. PLOS ONE 2018;13:e0207102.

24. Hu X, Yang Q. Real-time correction of image rotation with adaptive optics scanning light ophthalmoscopy. J Opt Soc Am A 2022;39:1663–1672.

25. Rodriguez AR, de Sevilla Müller LP, Brecha NC. The RNA binding protein RBPMS is a selective marker of ganglion cells in the mammalian retina. J Comp Neurol 2014;522:1411–1443.

26. Puthussery T, Gayet-Primo J, Taylor WR. Carbonic anhydrase-related protein VIII is expressed in rod bipolar cells and alters signaling at the rod bipolar to AII-amacrine cell synapse in the mammalian retina. Eur J Neurosci 2011;34:1419–1431.

27. Ma ICK, Nasir-Ahmad S, Lee SCS, et al. Contribution of parasol-magnocellular pathway ganglion cells to foveal retina in macaque monkey. Vision Res 2023;202:108154.

28. Peng Y-R, Shekhar K, Yan W, et al. Molecular Classification and Comparative Taxonomics of Foveal and Peripheral Cells in Primate Retina. Cell 2019;176:1222–1237.e22.

29. Dacey DM. The mosaic of midget ganglion cells in the human retina. J Neurosci 1993;13:5334–5355.

30. Tian L, Hires SA, Mao T, et al. Imaging neural activity in worms, flies and mice with improved GCaMP calcium indicators. Nature Methods 2009;6:875–881.

31. J. -S. Shyu, M. Maia, J. D. Weiland, et al. Electrical Stimulation in Isolated Rabbit Retina. IEEE Transactions on Neural Systems and Rehabilitation Engineering 2006;14:290–298.

32. Goo YS, Ye JH, Lee S, et al. Retinal ganglion cell responses to voltage and current stimulation in wild-type and rd1 mouse retinas. Journal of Neural Engineering 2011;8:035003.

33. Cho A, Ratliff C, Sampath A, Weiland J. Changes in ganglion cell physiology during retinal degeneration influence excitability by prosthetic electrodes. Journal of Neural Engineering 2016;13:025001.

34. Shiga Y, Rangel Olguin AG, Alarcon-Martinez L, et al. Light-evoked RGC calcium dynamics are altered in glaucoma: live imaging evidence of abnormal calcium clearance. Investigative Ophthalmology & Visual Science 2022;63:1850–1850.

35. Shiga Y, Rangel Olguin AG, El Hajji S, et al. Calcium clearance deficits linked to endoplasmic reticulum stress are signature features of early retinal ganglion cell damage in glaucoma. Investigative Ophthalmology & Visual Science 2023;64:482–482.

36. Gutierrez C, Cusick CG. Effects of chronic monocular enucleation on calcium binding proteins calbindin-D28k and parvalbumin in the lateral geniculate nucleus of adult rhesus monkeys. Brain Res 1994;651:300–310.

37. Sahel J-A, Boulanger-Scemama E, Pagot C, et al. Partial recovery of visual function in a blind patient after optogenetic therapy. Nature Medicine 2021;27:1223–1229.

