## Supplemental Figure 1 for "Foveal RGCs develop altered calcium dynamics weeks after photoreceptor ablation"

**Supp. 1: OCT data through region of PRL ablation in three subjects pre AAV injection, X weeks post AAV injection, and post PRL ablation.** (A) 30-degree OCT image through the region of high intensity ultrafast laser exposure in subject M1 43 weeks before AAV injection. (B) 30-degree OCT image through the two regions of sequential high intensity ultrafast laser exposure in subject M2 one week before AAV injection. (C) 30-degree OCT image through the region of high intensity ultrafast laser exposure in subject M3 72 weeks before AAV injection. (D) 30-degree OCT image through the region of high intensity ultrafast laser exposure in subject M1 1 week post AAV injection. (E) 30-degree OCT image through the two regions of sequential high intensity ultrafast laser exposure in subject M2 1 week post AAV injection. (F) 30-degree OCT image through the region of high intensity ultrafast laser exposure in subject M3 2 weeks post AAV injection. (G) 30-degree OCT image through the region of high intensity ultrafast laser exposure in subject M1, 119 weeks post AAV injection. (H) 30-degree OCT image through the two regions of sequential high intensity ultrafast laser exposure in subject M2, 7 weeks post AAV injection. Left: PRL ablation delivered 2 years before the study. Right: PRL ablation delivered 1 week before the study. (I) 30-degree OCT image through the region of high intensity ultrafast laser exposure in subject M3, 54 weeks post AAV injection.

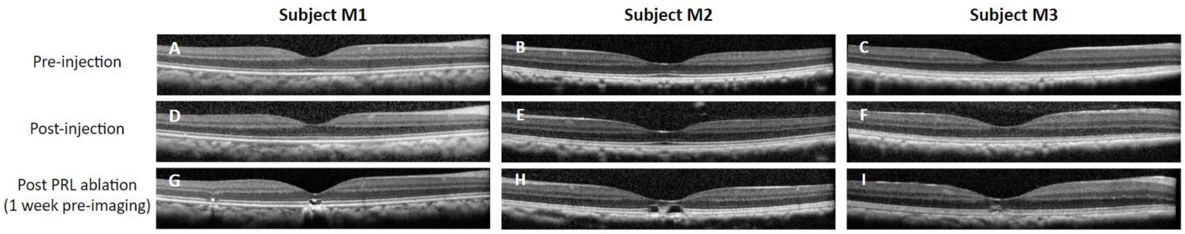
