## Supplemental Figure 2 for "Foveal RGCs develop altered calcium dynamics weeks after photoreceptor ablation"

**Supp. 2: Co-expression of viral vectors AAV2-CAG-GCaMP6s and AAV2-CAG-tdTomato:ChrimsonR in the foveal RGCS of the 3 NHPs involved in this study.**

(A) Fundus fluorescence image of GCaMP6s before PRL ablation in subject M1. (B) Fundus fluorescence image of GCaMP6s fluorescence before PRL ablation in subject M2. (C) Fundus fluorescence image of GCaMP6s before PRL ablation in subject M3. (D) Fundus fluorescence image of tdTomato before PRL ablation in subject M1. (E) Fundus fluorescence image of tdTomato before PRL ablation in subject M2. (A) Fundus fluorescence image of tdTomato before PRL ablation in subject M3.

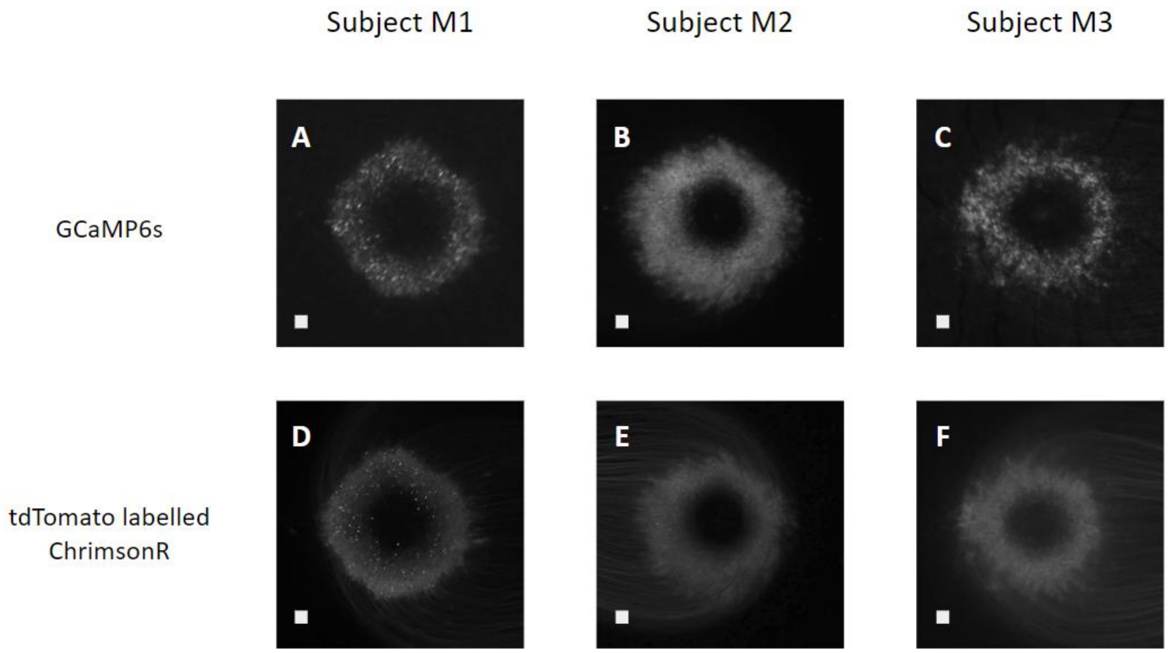
