## Supplemental Figure 3 for "Foveal RGCs develop altered calcium dynamics weeks after photoreceptor ablation"

**Supplemental Figure 3: Transfection of AAV2-CAG-GCaMP6s and AAV2-CAG-tdTomato:ChrimsonR in RGCs.** (A) Confocal image of a retinal section from the parafoveal region of animal M3 showing nuclei labeled with Hoescht in the outer nuclear layer (ONL), inner nuclear layer (INL) and ganglion cell layer (GCL). (B) Same retinal slice as in (A) showing immunolabeling of parasol RGCs with CAVIII (magenta) and endogenous GCaMP (green). The ROI in the GCL is shown enlarged in (C-E). Arrow shows an example of a parasol RGC transfected with GCaMP (E). On average, 2.2% of the GCaMP6s+ cells were also CAVIII+ (n=2 retinal sections). (F) Same retinal slice as in (A,B) showing immunolabeling of parasol RGCs with CAVIII (magenta) and endogenous tdTomato (green). The ROI in the GCL is shown enlarged in (G-I). Arrows show examples of parasol RGCs transfected with tdTomato. On average, 3.6% of the tdT+ cells were also CAVIII+ (n=2 retinal sections). Note that CAVIII also labels bipolar cells in the INL. Scale bars, 100 µm (A, B, F). Scale bars, 10 µm (C-E), (G-I).

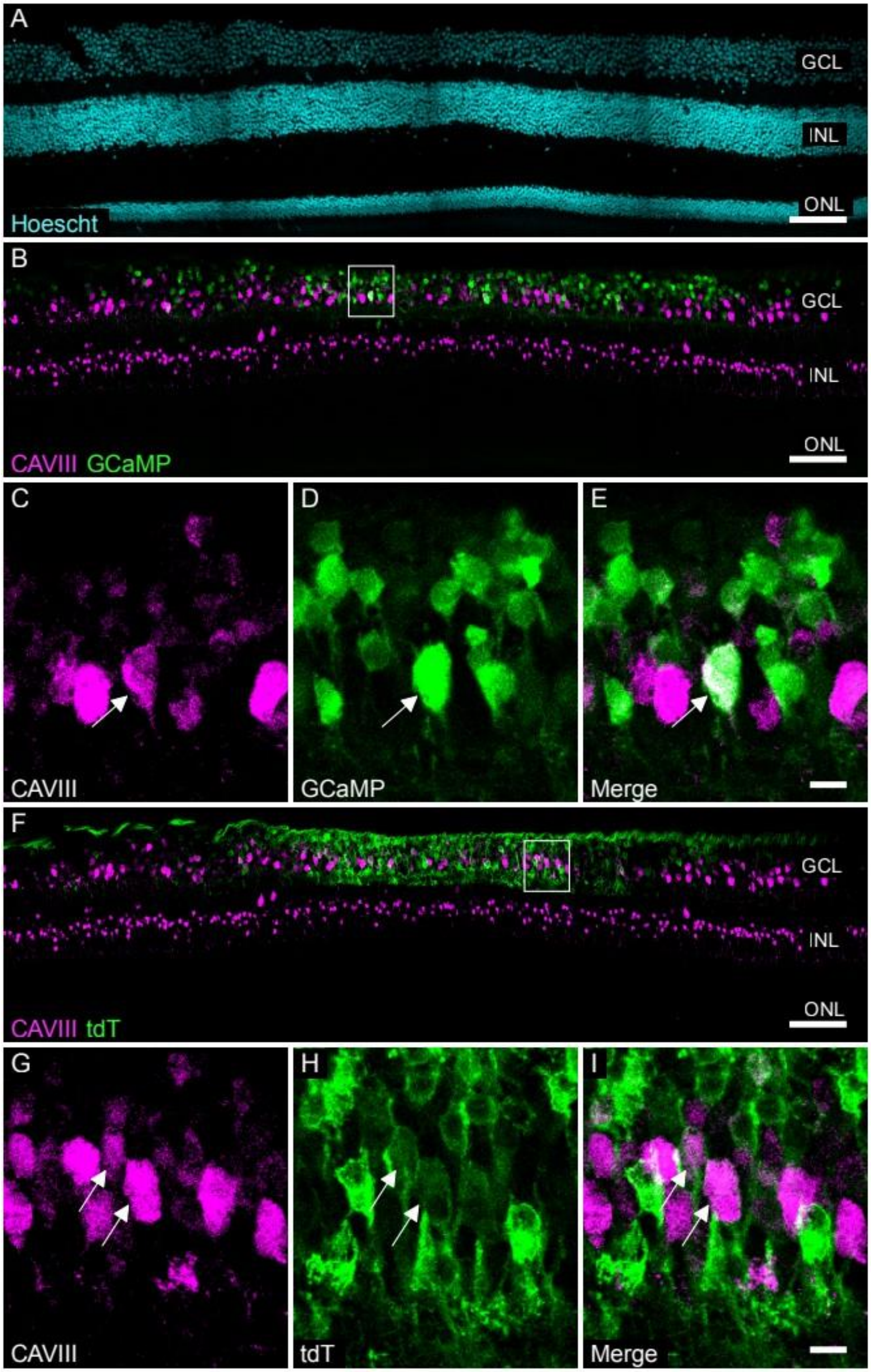
A
GCL
INL
ONL
Hoescht
B
GCL
INL
ONL
CAVIII GCaMP
C
CAVIII
D
GCaMP
E
Merge
F
GCL
INL
ONL
CAVIII tdT
G
CAVIII
H
tdT
I
Merge
