## Supplemental Figure 4 for "Foveal RGCs develop altered calcium dynamics weeks after photoreceptor ablation"

**Supp. 4: Localized Stimulation of RGCs with Intact Photoreceptors in primate foveal.** (A) Adaptive optics GCaMP 6s fluorescence image shows four locations of intact RGCs, and optogenetic stimulus is placed in the red square zone. Recordings were collected in the stimulated and nearby RGCs to show the exclusion of photoreceptor inputs from the localized stimulus. (B) Optogenetic responses of RGCs in stimulated and nearby locations, GCaMP fluorescence response were only present in the location of the stimulus, where nearby RGCs shows an absence of response to stimulus light.  All scale bars are 100 μm, and all data are acquired from subject M2.

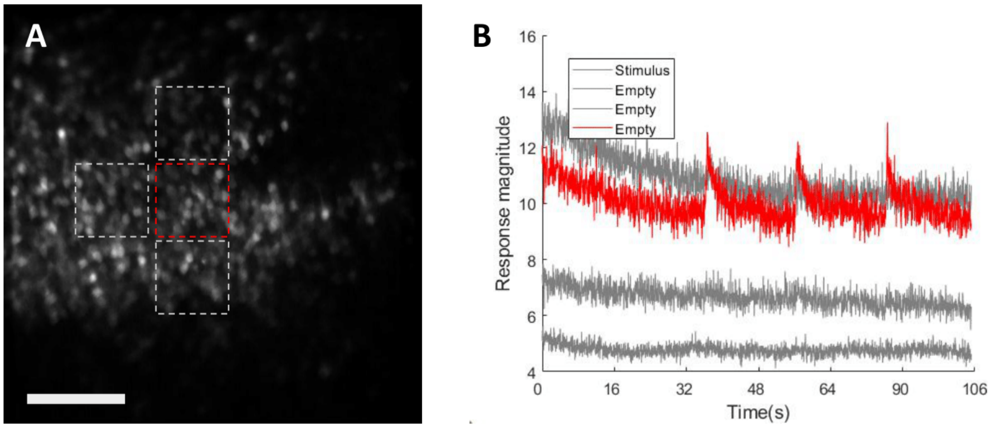
