## Supplemental Figure 5 for "Foveal RGCs develop altered calcium dynamics weeks after photoreceptor ablation"

**Supp. 5: Normalized response magnitude and normalized pre-stimulus baseline from short-term deafferented RGCs in the 10 weeks following photoreceptor ablation.** (A) Normalized optogenetic response magnitude from short-term deafferented RGCs in 10 weeks following photoreceptor ablation. (B) Normalized pre-stimulus baseline from short-term deafferented RGCs in 10 weeks following photoreceptor ablation. All Data collected from 109 short-term deafferented RGCs and 109 long-term deafferented RGCs in subject M2.

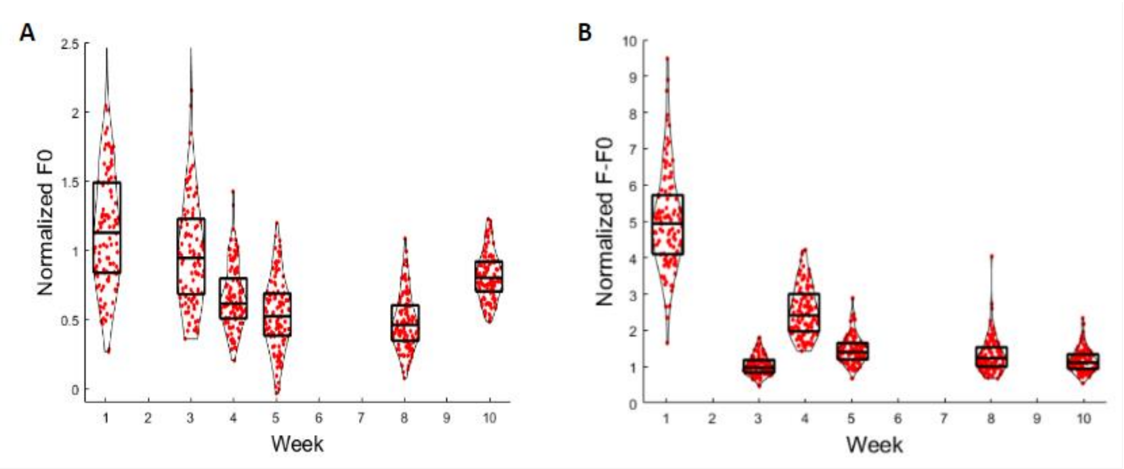
